# Longer Is Not Always Better: Effects of Equilibration Length on Umbrella Sampling Estimations for RNA Hairpin Folding Stabilities

**DOI:** 10.64898/2026.09.15.751614

**Authors:** Olayinka Akinyemi, Elzbieta Kierzek, Ryszard Kierzek, James P. McSally, Anees Mohammed Keedakkatt Puthenpeedikakkal, David H. Mathews

## Abstract

Umbrella sampling is widely used to estimate biomolecular free energy landscapes and relative folding stabilities. Although equilibration is a critical component of umbrella sampling workflows, the impact of equilibration length on thermodynamic predictions remains poorly understood. Here, we investigate the effect of equilibration length on relative folding free energy predictions for four RNA hairpins with loop sequences GUGAAA, CUGGGA, GUAAUA, and UUAAUU with helical stems of three base pairs. Umbrella sampling simulations were performed using an end-to-end distance reaction coordinate spanning 15-45 Å, where equilibrium simulations (windows) were spaced at roughly 1 Å intervals. In these calculations, the hairpin stem-loops were allowed to equilibrate in an end-to-end distance window and then the coordinates were transferred to the next larger end-to-end distance window to equilibrate. Two equilibration lengths, 2 ns and 100 ns per window, were followed by 600 ns of production sampling. Potential of mean force (PMF) profiles were reconstructed using the Weighted Histogram Analysis Method (WHAM) and used to calculate pairwise free energy differences with thermodynamic cycles. Increasing the equilibration length produced substantial, sequence-dependent changes in the reconstructed free energy landscapes. The 100 ns protocol generated markedly flatter PMFs for GUGAAA and UUAAUU and pronounced reshaping of the free energy landscape for GUAAUA. These changes were accompanied by reductions in hydrogen-bonding and stacking interactions, particularly within the intermediate regions of the reaction coordinate. The resulting thermodynamic predictions, as free energy change differences, were therefore highly sensitive to equilibration length. Across nearly all hairpin pairs, the 100 ns equilibration yielded substantially larger magnitude free energy change difference values than the corresponding 2 ns equilibration, with differences that greatly exceeded replica-to-replica variability. Comparison with optical melting measurements and nearest-neighbor thermodynamic predictions revealed that 2 ns equilibration times more closely agreed with experimental values than those obtained using 100 ns equilibration. These findings demonstrate that longer equilibration can systematically alter the structural ensembles sampled during umbrella sampling and amplify predicted stability differences without improving agreement with experiment. More widely, our results highlighted equilibration length as a critical and nontrivial parameter in RNA free energy calculations and demonstrate that increased equilibration does not necessarily lead to more accurate thermodynamic predictions.

## INTRODUCTION

Ribonucleic acid (RNA) plays important roles in numerous biological processes. Although earlier understood to primarily function in the expression of proteins, there is a wide variety of RNA, known as non-coding RNA, that are involved in a range of other processes. (1–3) RNA molecules play central roles in gene regulation, catalysis, molecular recognition, and cellular signaling. (4–6) These functions are intimately linked to RNA structure and dynamics, which are governed by a delicate balance of base pairing, base stacking, electrostatic interactions, and solvent effects. Even small changes in sequence can substantially alter the stability of RNA secondary structure motifs such as hairpin, internal, bulge, and multibranch loops. (7,8) Understanding the thermodynamics of RNA structural motifs remains an important objective of computational biophysics and is essential for understanding RNA function and for developing predictive models of RNA structure. (9–14)

Molecular dynamics (MD) simulations provide a powerful framework for investigating RNA conformational ensembles at atomic resolution. Advances in RNA force fields and computational resources have enabled increasingly realistic simulations of RNA folding, structural transitions, and ligand recognition processes. (15,16) However, many biologically relevant conformational changes occur on timescales that remain difficult to access using conventional MD simulations. As a result, enhanced-sampling methods have become indispensable tools for exploring RNA free-energy landscapes and estimating thermodynamic quantities such as folding free energies and relative stabilities. (17–20)

Among these approaches, umbrella sampling remains widely used for calculating free-energy profiles along predefined reaction coordinates. (21–24) In umbrella sampling, a series of biased simulations is performed to enhance sampling along a collective variable, and the resulting distributions are combined using methods such as the Weighted Histogram Analysis Method (WHAM) to reconstruct the underlying free energy landscape. (25–30) Umbrella sampling has been successfully applied to a wide range of RNA systems, including hairpin folding, base-pair disruption, tertiary structure formation, and ligand-induced conformational transitions.

The accuracy of umbrella sampling calculations depends critically on adequate sampling within individual equilibrium simulations, which we call windows. Considerable effort has therefore been devoted to selecting appropriate reaction coordinates to map out important biophysical processes. (31–33) In contrast, comparatively less attention has been paid to the role of equilibration time within umbrella windows. Equilibration is generally assumed to improve the quality of free energy estimates by allowing trajectories to relax from their initial configurations before data collection begins. Consequently, longer equilibration periods are assumed to be inherently beneficial for convergence. However, this hypothesis has rarely been examined systematically, particularly in RNA systems where multiple metastable conformational states may coexist at similar values of a chosen reaction coordinate.

This issue may be particularly important when the reaction coordinate does not uniquely define the underlying structural ensemble. For example, an RNA hairpin characterized by a particular end-to-end distance may still adopt a variety of conformations that differ substantially in hydrogen-bonding patterns, base-stacking interactions, loop geometry, and other structural features. If transitions among these substates occur on timescales comparable to or longer than the equilibration period, the resulting free energy estimates may depend strongly on the conformational ensemble sampled within each umbrella window. In such cases, increasing equilibration time may alter the populations of structurally distinct states without necessarily improving agreement with experimentally-determined free energy changes.

In prior work by Smith *et al*., we estimated the relative folding stability of hairpin stem-loops with sequence CGA<u>CAGUGC</u>UCG, GGC<u>GUAAUA</u>GCC and GCG<u>UUAAUU</u>CGC, where the underlined nucleotides are unpaired, using a combination of all-atom molecular dynamics simulation and umbrella sampling. We defined the reaction coordinate as the end-to-end distance of the hairpins, that is, the distance between 5’ and 3’ hydroxyl oxygens. The unbiased probabilities of conformations having a range of end-to-end distance from 15 Å to 60 Å were calculated. Relative folding stabilities were estimated from the sums of weighted probabilities corresponding to bins above and below a cuff-off distance of 45 Å. (34)

In this work, we investigate the effect of equilibration length on the estimation of relative folding stability of four RNA hairpin stem-loops: including new sequences GCG<u>GUGAAA</u>UGC (35) and GGC<u>CUGGGA</u>GCC (36) and previously studied sequences GGC<u>GUAAUA</u>GCC and GCG<u>UUAAUU</u>CGC using umbrella sampling molecular dynamics. We cover the range of end-to-end distances from 15 to 45 Å, with spacing between windows of 1 Å. In our setup, we equilibrate the hairpin stem-loop in the window, with a 15 Å end-to-end distance. Then we transfer the coordinates to the 16 Å window, equilibrate, and transfer to the next window, repeating up to 45 Å. Two equilibration times were used, 2 ns and 100 ns. After the equilibration period, each window is sampled for 600 ns. Reconstructed free energy landscapes were used to estimate relative folding free energies through thermodynamic cycles and these were compared with nearest-neighbor predictions and optical melting measurements. We report optical melting experiments for GCG<u>GUGAAA</u>UGC and GGC<u>CUGGGA</u>GCC, although GGC<u>CUGGGA</u>GCC did not have a melting transition as expected. To identify the structural origins of any equilibration-dependent effects in the simulations, we further analyzed hydrogen-bonding and base-stacking interactions along the unfolding pathway.

We hypothesized that the longer equilibration periods would result in more accurate free energy estimates, but our results suggest that increased equilibration generally worsened agreement with experiments. We found that equilibration length exerts a surprisingly strong influence on the reconstructed free energy landscapes and resulting thermodynamic predictions with more pronounced impact on UUAAUU and GUGAAA. Longer equilibration systematically altered the structural ensembles sampled within umbrella windows, modified hydrogen-bonding and stacking interactions, and overestimated the magnitude of estimated free energies change differences. These findings highlight equilibration length as a critical but often overlooked parameter in RNA free energy calculations and emphasize the importance of considering structural variables beyond the primary reaction coordinate in umbrella sampling simulations.

## RESULTS

### Hairpins Description

Four RNA hairpin stem-loops, each with a 6-nucleotide loop sequence and a three base pair stem were selected for this study. We refer to the hairpin stem-loops by their respective 6-nucleotide loop sequences. The starting structures are depicted in Figure 1.

**Figure 1:**
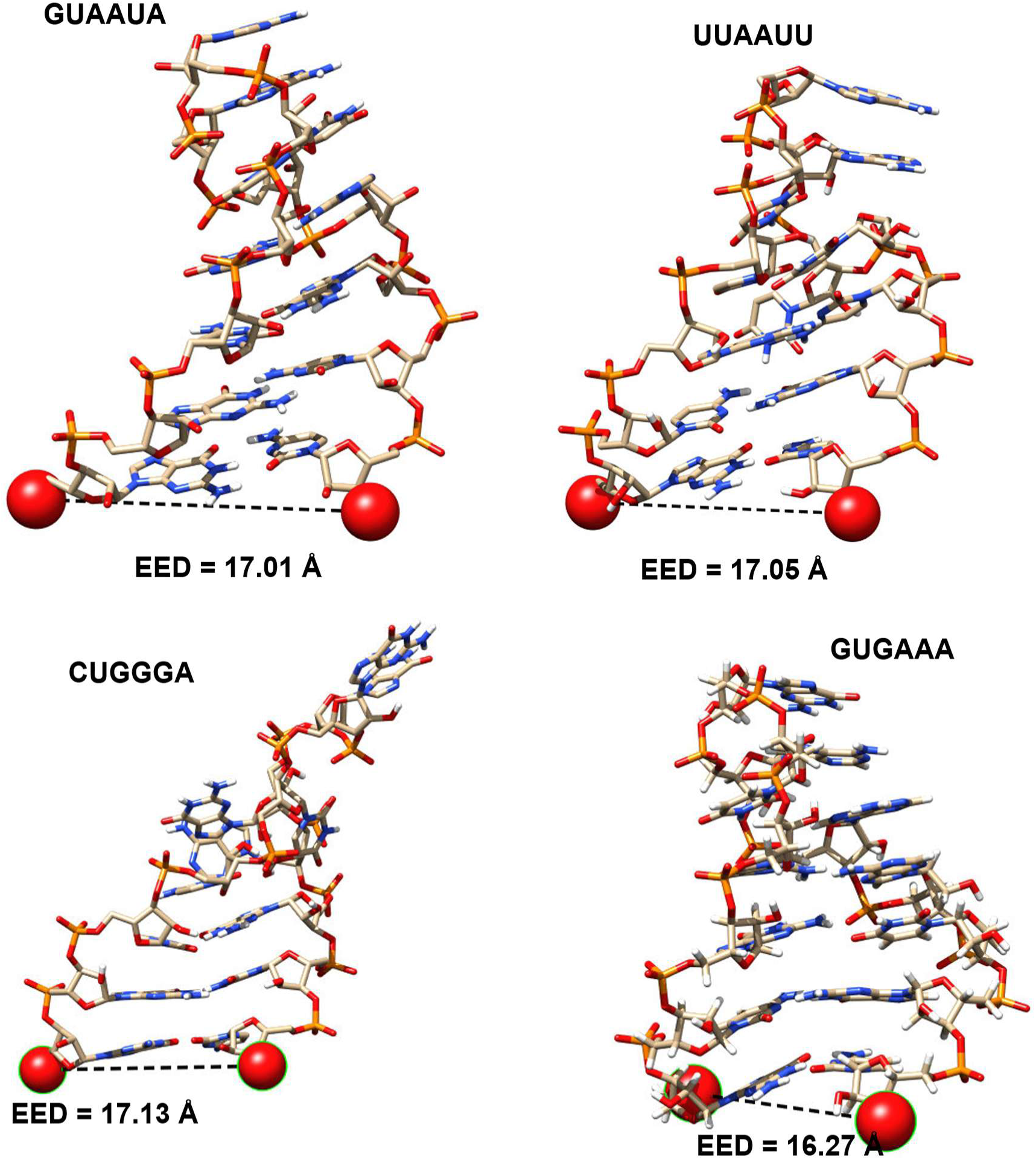
Initial structures used for umbrella sampling simulations of the four RNA hairpin stem-loops. Representative folded conformations of the hairpins with loop sequences GUAAUA, UUAAUU, CUGGGA, and GUGAAA used as starting structures for umbrella sampling. Red spheres indicate the terminal oxygen atoms used to define the end-to-end distance (EED) reaction coordinate, shown by the dashed line. The initial EED values correspond to the folded state and served as the starting point for the serial generation of umbrella windows spanning the unfolding pathway.

Two of the hairpins, GUAAUA and UUAAUU are described in and previously studied by Smith *et al.* and were reported to have a relative folding free energy, ΔΔG°stretch, of 2.58 + 0.44 kcal/mol. (34) GUAAUA was also reported in Spasic *et al.*, where the free energy change of helix formation was studied. (37) Both sets of starting coordinates were taken from solution structures (PDB ID: 1HS3). (38)

The remaining two hairpins, GUGAAA and CUGGGA, were available as NMR structures. Neither had been previously studied by optical melting experiments or by umbrella sampling. GUGAAA is the structure of the 690 loop of the *E. coli* small subunit rRNA (PDB ID: 1FHK) sourced from the protein databank (PDB) with its terminal base pair excised, leaving a 12-nucleotide sequence (GCG<u>GUGAAA</u>UGC). (35) This truncation to ten nucleotides is performed to match the length of the previously studied hairpin stem-loops. CUGGGA is the structure of the HIV-2 TAR-Argininamide complex (PDB ID: 1AJU). (36) A terminal AU pair in the stem was replaced by a GC base pair to improve the stability of the stem, resulting in sequence GGC<u>CUGGGA</u>GCC.

### Longer Equilibration Produces Sequence-Dependent Changes in RNA Free Energy Landscapes

To assess the influence of equilibration length on free energy estimation, potential of mean force (PMF) profiles were reconstructed from umbrella sampling simulations performed using 2 ns and 100 ns equilibration protocols (Figure 2). To determine the sampling uncertainty, four replicas of each system were simulated. For all four RNA hairpins, the PMFs exhibit a global minimum at ∼17 Å corresponding to folded conformations and increase as the molecules are progressively extended. However, substantial differences in PMF shape and replica behavior are observed between the two equilibration schemes. The strongest effect of equilibration length is observed for the GUGAAA and UUAAUU hairpins. Compared to the 2 ns equilibration simulations, the corresponding 100 ns PMFs generally display a substantially flatter free energy increase across the reaction coordinate. In both systems, the free energy penalty associated with extension is reduced over a wide range of end-to-end distances, resulting in less steep transitions from folded to extended conformations. The GUAAUA hairpin exhibits a different response to equilibration length. Although the 100 ns PMF differs substantially from the 2 ns profile, the dominant feature is not a simple flattening of the free energy landscape. Instead, the PMF undergoes noticeable reshaping, particularly in the intermediate end-to-end distance region (∼28-35 Å). This behavior is accompanied by increased variability among independent replicas in the 100 ns equilibration simulations relative to the 2 ns protocol, indicating a greater sensitivity of the sampled free energy landscape to equilibration conditions. Among the four systems, CUGGGA displays the weakest dependence on equilibration length. While modest broadening of the folded basin is observed following 100 ns equilibration, the overall shape of the PMF remains qualitatively similar to that obtained using the shorter equilibration protocol.

**Figure 2.**
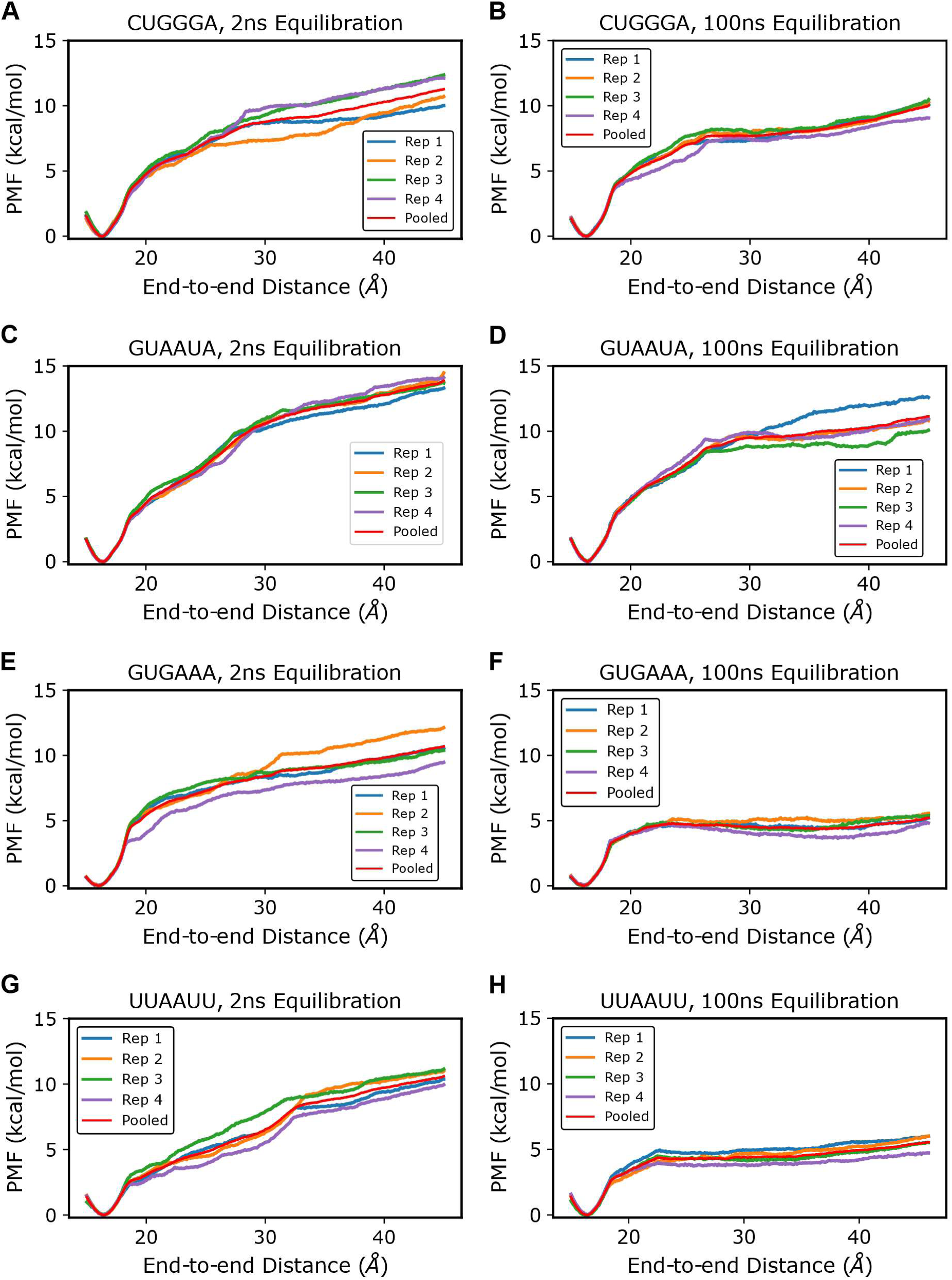
Replica and pooled PMFs for the four RNA hairpins. Potential of mean force (PMF) profiles reconstructed from four independent umbrella sampling replicas (Rep 1–4) and pooled trajectories (purple) for the CUGGGA, UUAAUU, GUAAUA, and GUGAAA hairpins. Simulations were performed using 2 ns (left column) and 100 ns (right column) equilibration periods followed by 600 ns of production sampling per umbrella window. PMFs were reconstructed using WHAM along the end-to-end distance (EED) reaction coordinate. The pooled PMFs were generated by combining trajectories from all four replicas and were subsequently used for relative folding free energy calculations.

The influence of equilibration length on replica-to-replica consistency is sequence dependent. For CUGGGA, GUGAAA, and UUAAUU, PMFs generated using 100 ns equilibration exhibit reduced spread among independent replicas relative to the corresponding 2 ns simulations, indicating greater reproducibility of the reconstructed free energy profiles. In contrast, GUAAUA shows the opposite trend, with tighter clustering of replica PMFs following 2 ns equilibration and increased divergence with 100 ns equilibration. These observations demonstrate that equilibration length influences both the shape and reproducibility of reconstructed free energy landscapes, although the magnitude and nature of these effects vary substantially between sequences.

### Predicted Relative Folding Free Energies Increase with Equilibration Length

Using the reconstructed PMFs, the folding free energy (ΔG°) was estimated at a cutoff end-to-end distance of 44 Å between slack and stretched states (Table 1). At this distance, the mean non-neighboring base-base hydrogen bond count fell below 0.3, at which point the systems can be regarded as fully stretched. ΔG° values were markedly lower in the shorter equilibration than the long scheme for GUGAAA (−6.91 ± 0.14 compared to −10.81 ± 0.46 kcal/mol) and UUAAUU (−5.72 ± 0.26 compared to −10.70 ± 0.24 kcal/mol) corresponding to the flattening observed in the PMFs with the longer equilibration scheme.

**Table 1.**
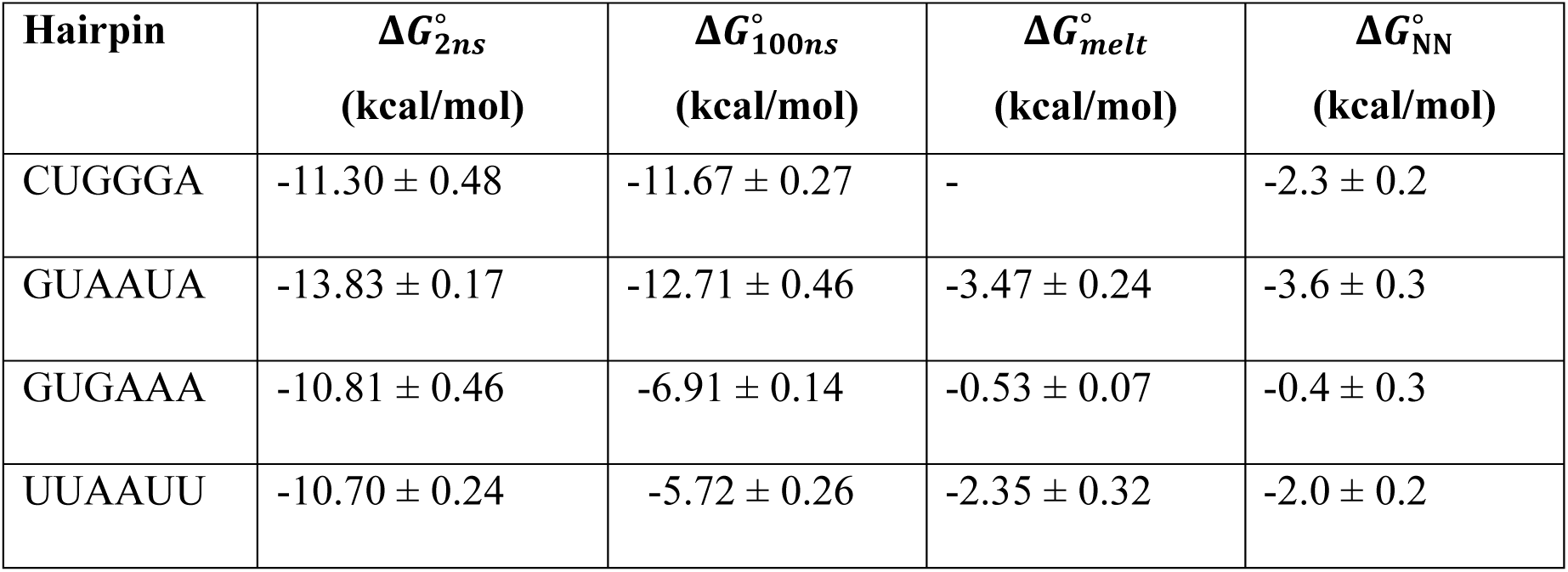
Folding free energies (Δ*G*°) at 37 °C of the RNA hairpins obtained from umbrella sampling simulations and experimental and nearest-neighbor predictions. Folding free energies were calculated from reconstructed PMFs with a cutoff end-to-end distance of 44 Å. Simulated values are reported for the 2 ns and 100 ns equilibration protocols, with uncertainties estimated as standard error of the mean (pooled simulation) of the 4 replicates. Experimental folding free energies (Δ*G*°) were obtained from optical melting measurements, while Δ*G*° values were calculated using nearest-neighbor thermodynamic parameters. (2,3) More negative values correspond to greater thermodynamic stability of the folded hairpin. Missing entries indicate that corresponding experimental measurements or nearest-neighbor estimates were unavailable.

Because of the markedly different denaturation free energy changes between the two equilibration schemes, we also performed optical melting experiments on GUGAAA and CUGGGA to compare to experiments (Table 1). CUGGGA did not demonstrate the expected shape for the denaturation of a hairpin, instead it showed a broad increase in UV absorption as a function of temperature. We expect this was in part caused by the 3 adjacent Guanines, which tend to lower strand solubility possibility because of G quadruplex formation (39–41). GUGAAA had a ΔG°37 of −0.53 ± 0.67 kcal/mol, ΔH° of −34.55 ± 1.47 kcal/mol, and ΔS° of −109.68 ± 4.92 cal/K·mol.

In the absence of an experimentally-determined stability for CUGGGA, we make comparisons instead to ΔG°37 estimates from nearest neighbor estimates. (42–44) Hairpin stem-loop stabilities for loops of six unpaired nucleotides are expected to be within 0.5 kcal/mol of the experimental value; the uncertainty in the initiation term for these loops is 0.5 kcal/mol and the RMSD between experiment and estimate for the other three hairpin stem-loops in this study is 0.3 kcal/mol (Table 1). (45)

The free energy changes estimated by simulation and those estimated by optical melting or nearest neighbor analysis are not directly comparable. The unpaired state in simulations is a stretched state that holds the ends far apart, which is much lower in entropy than the random coil state from optical melting and nearest neighbor analysis. (37,46,47) Relative folding free energies (ΔΔG°), however, are comparable and these were calculated for all pairwise combinations of the four RNA hairpins using a thermodynamic cycle (Figure 3). The resulting ΔΔG° values exhibit a strong dependence on equilibration length (Table 2). Notably, the direction of relative folding stability was correctly predicted as compared to optical melting or nearest neighbor analysis by MD simulation for all pairs of sequences except for GUGAAA-UUAAUU, although the longer equilibration scheme was farther off (1.2 ± 0.3 kcal/mol) than the shorter (0.12 ± 0.52).

**Figure 3.**
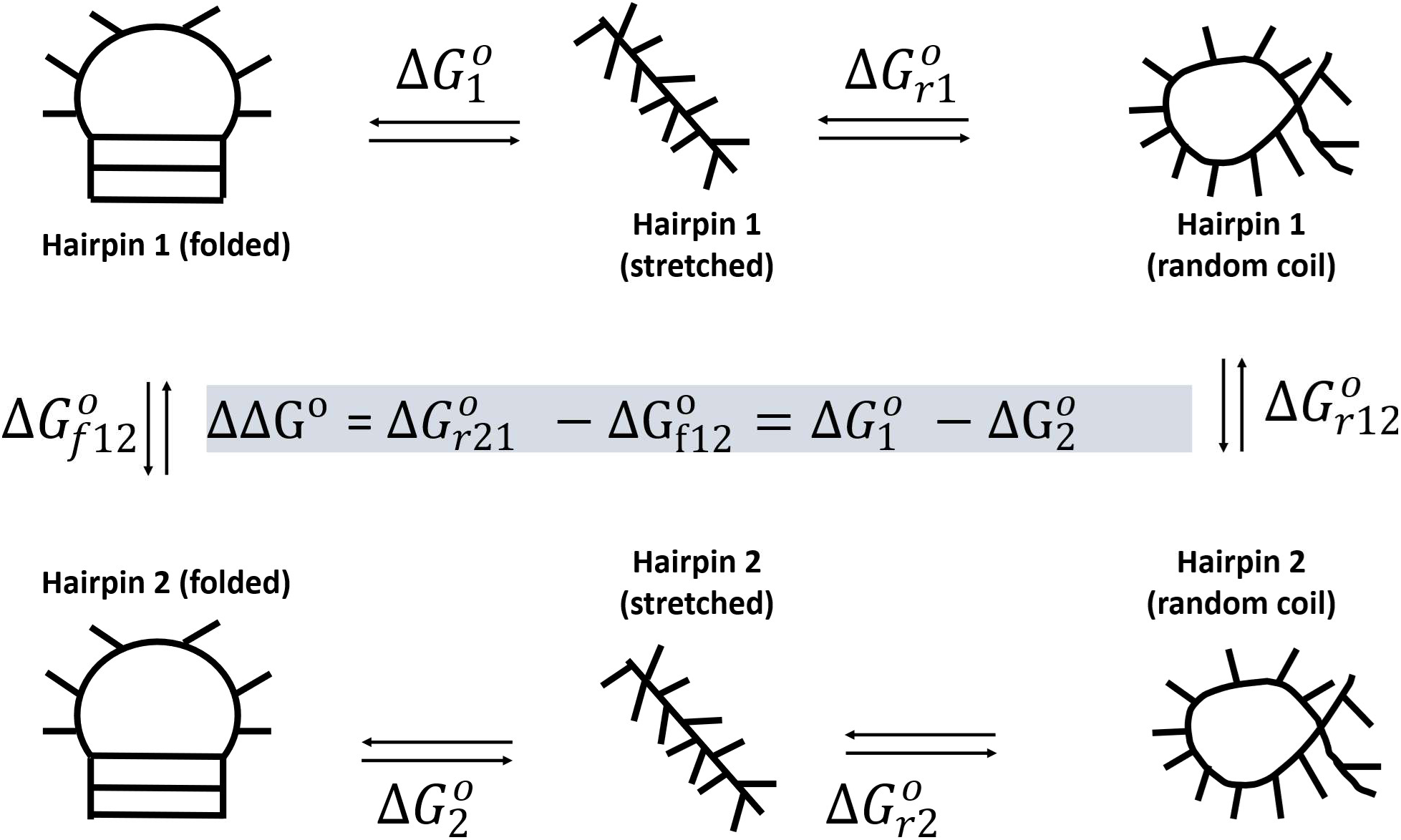
Thermodynamic cycle relating simulated and experimental folding free energies. Folding free energies were decomposed into folded-to-stretched and stretched-to-random-coil contributions. Umbrella sampling was used to estimate the folded-to-stretched free energies (Δ*G*°_1_ and Δ*G*°_2_) for each hairpin. Assuming that the stretched-to-random-coil relaxation free energy (Δ*G*°_r2_ *and* Δ*G*°_r1_) is sequence independent, (1) these contributions cancel in the thermodynamic cycle, yielding ΔΔ*G*° = Δ*G*°_1_ − Δ*G*°_2_ = Δ*G*°_ƒ12_ − Δ*G*°_r12_, where the difference in the vertical legs is the difference between two optical melting experimentsThe resulting ΔΔ*G*°values represent relative folding stabilities and were compared with optical melting experiments and nearest-neighbor thermodynamic predictions.

**Table 2.** Relative folding free energies (ΔΔ*G*°) obtained from umbrella sampling simulations, optical melting and nearest-neighbor predictions. Relative folding free energies were calculated using the thermodynamic cycle shown in Figure 3. Experimental values (ΔΔ*G*°_2ns_) were derived from optical melting measurements, while ΔΔ*G*°_100ns_ values were calculated from nearest-neighbor thermodynamic parameters. ΔΔ*G*° and ΔΔ*G*° are predicted values from 2 ns and 100 ns equilibration simulations, respectively. Negative values indicate that the product hairpin is predicted to be more stable than the reactant, whereas positive values indicate greater stability of the reactant hairpin. Missing entries indicate that corresponding experimental measurements were not available.

| Reactant | Product | $\Delta\Delta G_{2\text{ns}}^\circ$ | $\Delta\Delta G_{100\text{ns}}^\circ$ | $\Delta\Delta G_{\text{NN}}^\circ$ | $\Delta\Delta G_{\text{melt}}^\circ$ |
| --- | --- | --- | --- | --- | --- |
| CUGGGA | GUAAUA | $-2.53 \pm 0.51$ | $-1.04 \pm 0.53$ | $-1.3 \pm 0.36$ | - |
| GUGAAA | CUGGGA | $-0.49 \pm 0.66$ | $-4.75 \pm 0.30$ | $-1.9 \pm 0.36$ | - |
| GUGAAA | GUAAUA | $-3.02 \pm 0.49$ | $-5.80 \pm 0.48$ | $-3.2 \pm 0.42$ | $-2.94 \pm 0.25$ |
| GUGAAA | UUAUUU | $0.12 \pm 0.52$ | $1.20 \pm 0.30$ | $-1.6 \pm 0.36$ | $-1.82 \pm 0.33$ |
| UUAUUU | CUGGGA | $-0.61 \pm 0.54$ | $-5.95 \pm 0.38$ | $-0.3 \pm 0.28$ | - |
| UUAUUU | GUAAUA | $-3.13 \pm 0.29$ | $-7.00 \pm 0.53$ | $-1.6 \pm 0.36$ | $-1.12 \pm 0.40$ |

Across nearly all hairpin stem-loop pairs, the 100 ns equilibration protocol produces substantially larger magnitude free energy differences than the corresponding 2 ns protocol. The largest differences are observed for system pairs involving GUGAAA and UUAAUU. For example, the ΔΔG° between CUGGGA and UUAAUU changes from −0.61 ± 0.54 kcal/mol in the 2 ns simulations to −5.95 ± 0.38 kcal/mol following 100 ns equilibration, while the ΔΔG° between CUGGGA and GUGAAA changes from −0.49 ± 0.66 kcal/mol to −4.75 ± 0.30 kcal/mol. Similar increases in magnitude are observed for GUAAUA and GUGAAA (−3.02 ± 0.49 kcal/mol to −5.80± 0.48 kcal/mol) and GUAAUA and UUAAUU (−3.13 ± 0.29 kcal/mol to −7 ± 0.53 kcal/mol). In contrast, the relative ordering of system stabilities remains qualitatively similar between equilibration protocols, with GUAAUA consistently predicted to be the most stable of the four hairpins. The magnitude of the equilibration-dependent changes in ΔΔG° greatly exceeds the replica-to-replica variability observed within either protocol, indicating that equilibration length represents a significant source of variation in the predicted thermodynamic stabilities. These results demonstrate that equilibration-dependent differences in the sampled conformational ensembles propagate directly into the reconstructed PMFs and consequently into the calculated relative folding free energies.

### Longer Equilibration Systematically Worsened Agreement with Reference Thermodynamics

To evaluate the thermodynamic accuracy of the umbrella sampling protocols, the calculated ΔΔG° values were compared with available optical melting measurements and nearest-neighbor (NN) thermodynamic predictions using Δ(ΔΔG°) (Table 3). Optical melting data were available for three hairpin pairs. In all three cases, the ΔΔG° values obtained using the 2 ns equilibration protocol were closer to experiment than those obtained using 100 ns equilibration. This trend is reflected in the root-mean-square deviation (RMSD) between simulated and reference relative folding free energies. Relative to optical melting measurements, the RMSD increased from 1.65 ± 0.34 kcal/mol for the 2 ns protocol to 4.00 ± 0.36 kcal/mol for the 100 ns protocol. Similarly, relative to nearest-neighbor predictions, the RMSD increased from 1.22 ± 0.27 kcal/mol to 3.74 ± 0.23 kcal/mol. These results demonstrate that longer equilibration systematically worsened agreement with both experimental and nearest-neighbor thermodynamic benchmarks.

**Table 3.**
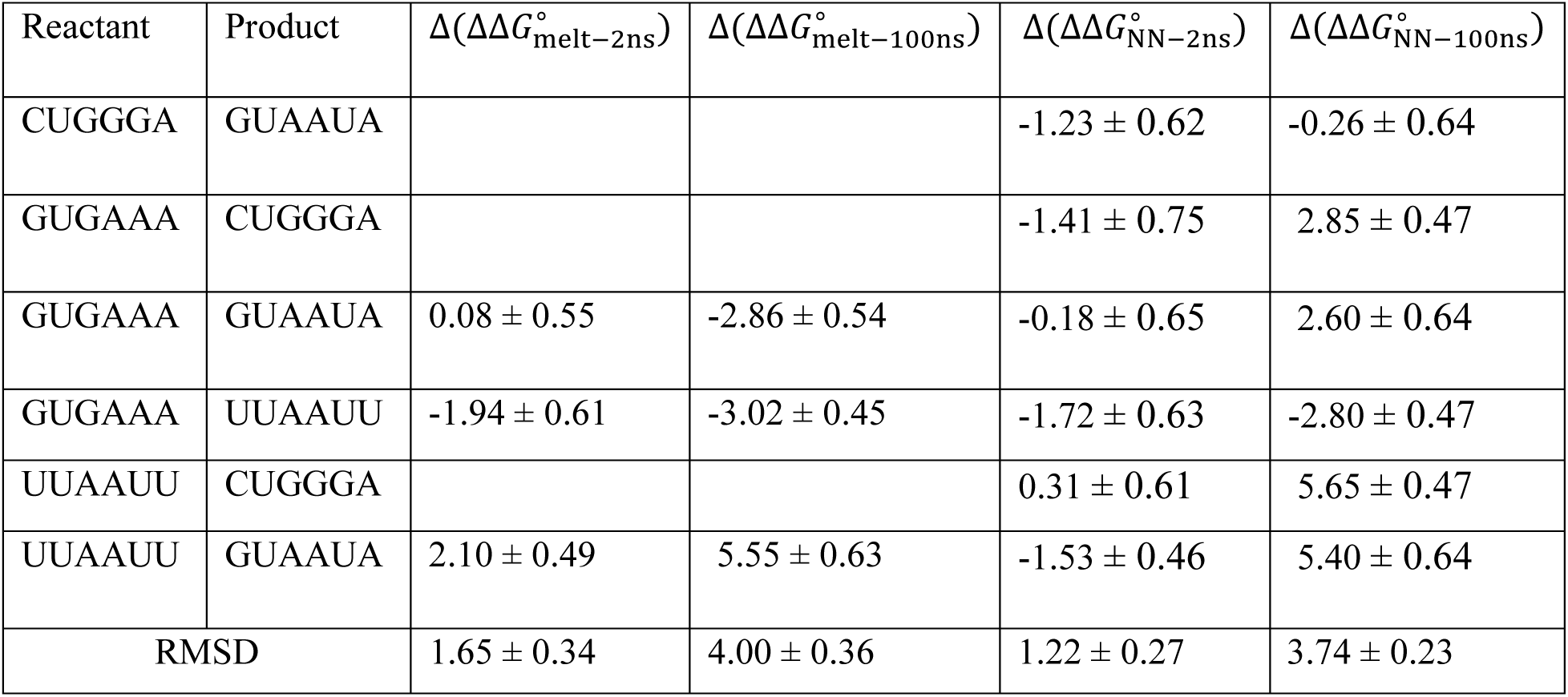
Deviation of simulated relative folding free energies from experimental and nearest-neighbor predictions. The quantities Δ(ΔΔ*G*°_melt–2ns_) and Δ(ΔΔ*G*°_melt–100n_) represent the differences between simulated and experimentally derived relative folding free energies, for the 2 ns and 100 ns equilibration protocols, respectively. Similarly, Δ(ΔΔ*G*°_NN–2n_) and Δ(ΔΔ*G*°_NN–100ns_) denote the differences between simulated and nearest-neighbor (NN) relative folding free energies. Values closer to zero indicate better agreement between simulation and the corresponding reference. Positive and negative values indicate overestimation and underestimation of the relative folding free energy difference, respectively. Uncertainties were obtained by propagating the errors associated with the simulated ΔΔ*G*° values. Blank entries indicate that the corresponding experimental measurements were unavailable.

The closest agreement was observed for the GUAAUA-GUGAAA pair, for which the 2 ns simulations predicted a ΔΔG° of −3.02 ± 0.49 kcal/mol compared with an experimental value of −2.94 ± 0.25 kcal/mol, corresponding to a deviation of only 0.08 ± 0.55 kcal/mol. In contrast, the corresponding 100 ns simulation yielded a ΔΔG° of −5.80 ± 0.48 kcal/mol, deviating from experiment by 2.86 ± 0.54 kcal/mol. Similar trends were observed for the UUAAUU-GUGAAA and GUAAUA-UUAAUU comparisons, where increasing the equilibration length resulted in substantially larger deviations from experiment.

The GUAAUA-UUAAUU Δ(ΔΔG°) was calculated as part of our previous study. In that work, the difference was 1.47 ± 0.57 kcal/mol, using a 2 ns equilibration period, but with TIP3P water. In this work, using OPC water, the 2 ns equilibration simulations estimated a Δ(ΔΔG°) of 2.10 ± 0.49 kcal/mol. This agreement is within the uncertainties between the two sets of simulations.

Comparison with NN thermodynamic predictions yielded a similar outcome. For five of the six pairwise comparisons, the 2 ns equilibration protocol produced ΔΔG° values that were closer to NN predictions than those obtained from the 100 ns equilibration simulations with the former resulting in an absolute difference as low of 0.18 ± 0.65 kcal/mol (GUAAUA-GUGAAA), with the maximum being 1.72 ± 0.63 kcal/mol (UUAAUU-GUGAAA). In contrast, the 100 ns equilibration simulations deviated in magnitude from NN results up to 5.65 ± 0.47 kcal/mol in CUGGGA-UUAAUU. The only exception was the GUAAUA-CUGGGA pair, for which the 100 ns result exhibited a smaller (0.26 kcal/mol + 0.64 compared to 1.23 + 0.62 kcal/mol) absolute deviation from the NN estimate.

Overall, these results demonstrate that equilibration length exerts a strong influence on the thermodynamic predictions derived from umbrella sampling simulations. Longer equilibration does not improve agreement with either optical melting experiments or nearest-neighbor (NN) thermodynamic predictions. Instead, the 100 ns equilibration protocol systematically amplifies the magnitude of the predicted stability differences between hairpins, leading to poorer agreement with available reference data.

### Hydrogen-Bond Networks Are Weakened by Extended Equilibration

To investigate the structural origin of the equilibration-dependent free energy landscapes, we quantified native and non-neighboring base-base hydrogen bonds as a function of end-to-end distance (Figure 4). For all four hairpins, the mean hydrogen-bond counts decrease as the molecules are extended, consistent with progressive disruption of folded structure during unfolding. However, substantial differences are observed between the 2 ns and 100 ns equilibration protocols.

**Figure 4.**
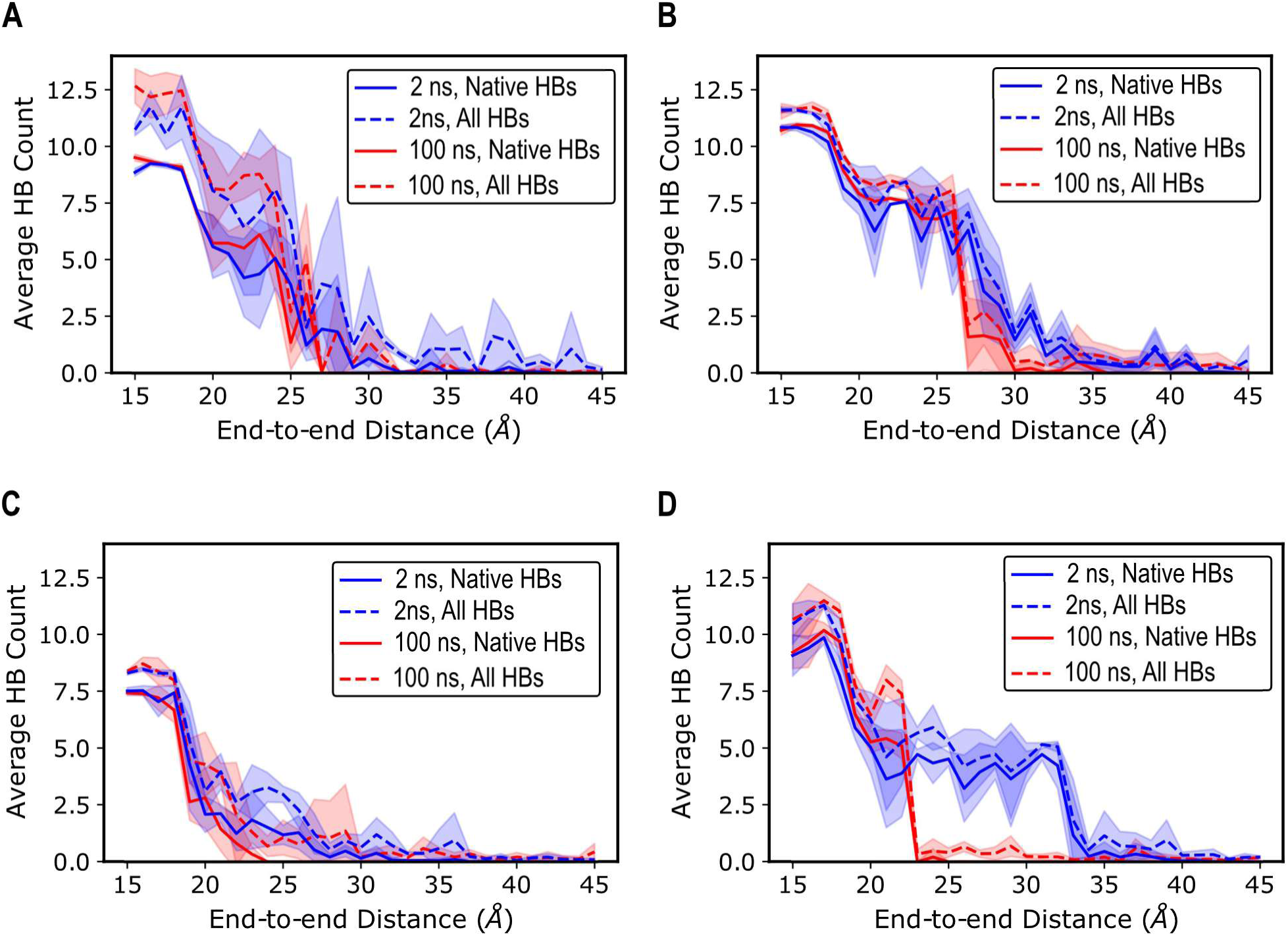
Hydrogen-bonding interactions along the unfolding coordinate. Mean hydrogen-bond counts as a function of end-to-end distance (EED) for the (A) CUGGGA, (B) GUAAUA, (C) GUGAAA, and (D) UUAAUU hairpins. Solid lines represent native hydrogen bonds present in the folded structure, while dashed lines represent all non-neighboring base-base hydrogen bonds. Blue and red curves correspond to the 2 ns and 100 ns equilibration protocols, respectively. Shaded envelopes indicate one standard deviation across four independent replicas.

The most pronounced effect is observed in the intermediate end-to-end distance regime, where partially folded conformations are expected to contribute significantly to the free energy landscape. In GUGAAA and UUAAUU, the 2 ns equilibration simulations retain substantially larger numbers of both native and non-neighboring base-base hydrogen bonds over a broad range of intermediate distances than the corresponding 100 ns simulations. This difference is particularly striking for UUAAUU, where the 2 ns simulations maintain approximately four to five hydrogen bonds for end-to-end distances between 24 and 32 Å, whereas the 100 ns simulations exhibit near-complete loss of hydrogen bonding over the same region.

Similar, although less dramatic, behavior is observed for GUGAAA and CUGGGA. GUAAUA exhibits a distinct pattern. At low end-to-end distances, both equilibration protocols sample similar hydrogen-bonding networks. However, as the hairpin is extended, the 2 ns and 100 ns simulations diverge, with the shorter equilibration protocol generally maintaining a larger number of hydrogen bonds throughout the transition region. Unlike the other systems, the differences between native and non-neighboring hydrogen-bond populations are comparatively small, suggesting substantial interconversion between alternative hydrogen-bonding arrangements during extension. Across all four systems, non-neighboring hydrogen bonds closely mirror the behavior of native hydrogen bonds rather than increasing as native contacts are lost. This observation indicates that disruption of the native stem is not accompanied by extensive formation of alternative non-native hydrogen-bonding networks. Instead, both native and non-neighboring hydrogen-bond populations generally decrease together as the hairpins unfold.

Taken together, these results demonstrate that equilibration length substantially influences the hydrogen-bonding ensembles sampled within umbrella windows. In most systems, the 100 ns equilibration protocol produces conformational ensembles with fewer hydrogen bonds in the intermediate regions of the reaction coordinate. These structural differences are consistent with the equilibration-dependent changes observed in the reconstructed free energy landscapes.

### Extended Equilibration Reduces Base Stacking During Hairpin Unfolding

To further investigate the structural origin of the equilibration-dependence of the PMFs, we quantified base stacking interactions along the end-to-end distance reaction coordinate for both equilibration protocols (Figure 5). For all systems, stacking interactions decreased progressively as the hairpins were extended, consistent with disruption of folded structure during denaturation with increasing end-to-end distance. However, clear differences were observed for some sequences between the 2 ns and 100 ns equilibration simulations, indicating that equilibration length substabtially influences the structural ensembles sampled within umbrella windows.

**Figure 5.**
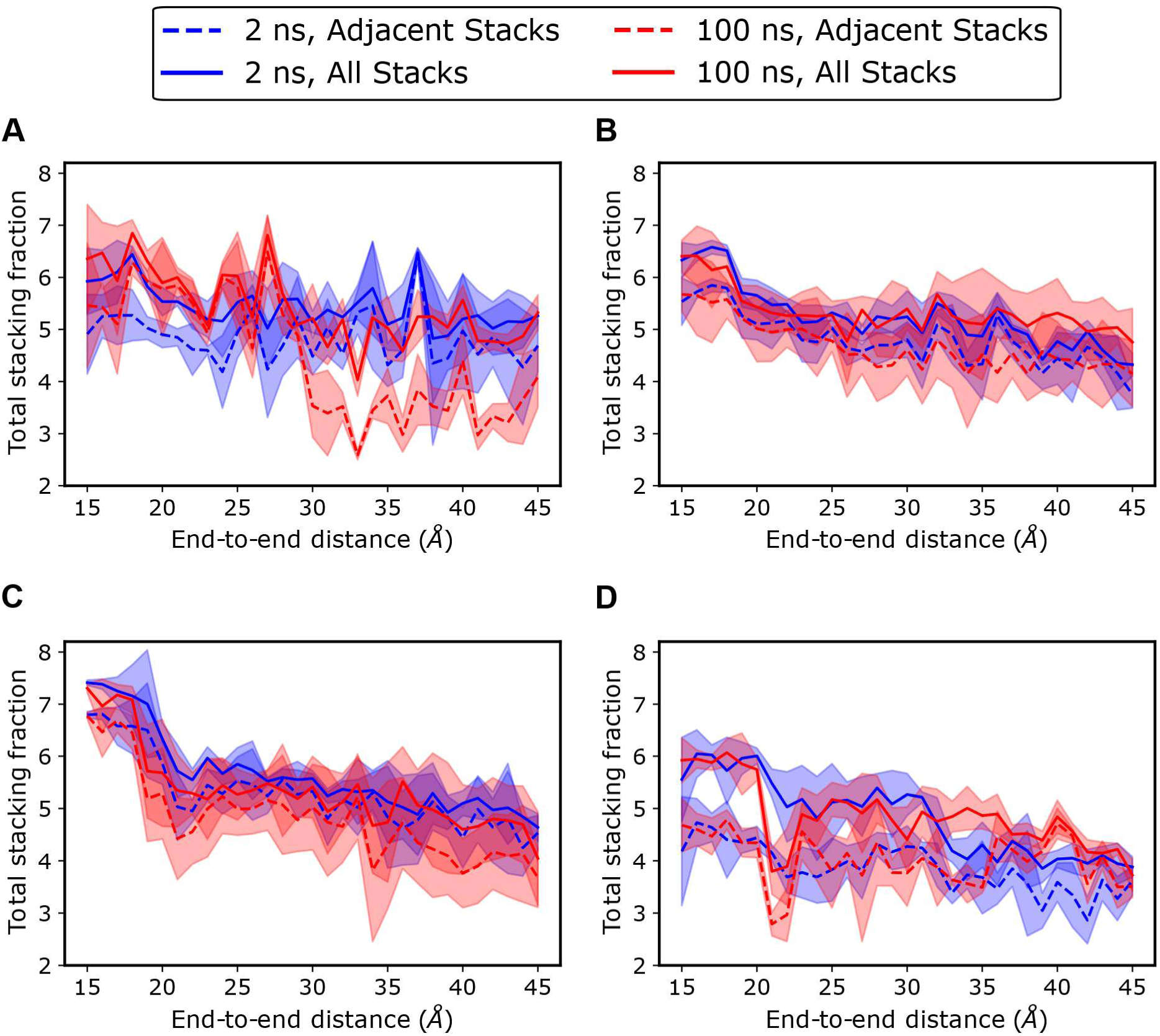
Base-stacking occupancies for the four RNA hairpins. Mean stacking occupancies as a function of end-to-end distance (EED) for the CUGGGA (A), GUAAUA (B), GUGAAA (C), and UUAAUU (D) hairpins. Solid lines denote total stacking occupancies summed over all residue pairs, while dashed lines denote occupancies summed over adjacent residue pairs only. Blue and red curves correspond to the 2 ns and 100 ns equilibration protocols, respectively. Shaded envelopes represent one standard deviation calculated from four independent replicas.

Like with hydrogen bonding, the most pronounced effects were observed for GUGAAA and UUAAUU. In both systems, the 2 ns equilibration simulations exhibited higher stacking persistence across a broad range of end-to-end distances, particularly when all stacking interactions were considered. By contrast, the 100 ns equilibration simulations showed reduced stacking fractions and an abrupt decay of stacking interactions with extension particularly in the intermediate region (∼20-35 Å) of UUAAUU. GUAAUA exhibited a different pattern of behavior. Although differences between equilibration protocols were still observed, the 2 ns and 100 ns equilibration profiles remained highly similar across the reaction coordinate, with only modest increases in stacking observed for the 100 ns simulations in the stretched region. CUGGGA showed a somewhat unique trend with greater stacking accompanying the 100 ns equilibration in the shorter end-to-end regions where adjacent stacks closely track the all-inclusive stacks, and a dramatic loss of adjacent stacking in the longer end-to-end regions.

In all systems, stacking interactions computed using all residue pairs were generally larger than those restricted to adjacent residue pairs, indicating the presence of non-neighboring stacking interactions throughout the unfolding process. Overall, the stacking analysis demonstrates that equilibration length alters the persistence and distribution of stacking interactions sampled during umbrella sampling simulations.

## DISCUSSION

Equilibration is generally regarded as a prerequisite for obtaining reliable free energy estimates from umbrella sampling simulations. A common assumption is that longer equilibration should improve convergence and therefore lead to more accurate thermodynamic predictions. Surprisingly, our results do not support this expectation. Across four RNA hairpins, extending the equilibration period from 2 to 100 ns produced substantial changes in the reconstructed free energy landscapes and systematically amplified the magnitude of the predicted relative folding free energies. More importantly, the resulting ΔΔG° values exhibited poorer agreement with both optical melting measurements and nearest-neighbor thermodynamic predictions.

The effects of equilibration length were evident at multiple levels of analysis. The 100 ns equilibration protocol produced pronounced flattening of the PMFs for GUGAAA and UUAAUU and substantial reshaping of the free energy landscape for GUAAUA (Figure 2). These changes were a result of reductions in hydrogen-bonding and stacking interactions throughout the unfolding coordinate compared to the shorter equilibration protocol, particularly within the intermediate end-to-end distance regime (Figures 4 and 5). The consistency of these observations suggests that the equilibration-dependent differences in ΔΔG° do not arise from statistical noise in the PMF reconstruction but instead reflect genuine differences in the structural ensembles sampled during umbrella sampling.

A natural question is why such structural differences should matter when the free energy calculations are performed along the end-to-end distance reaction coordinate. In principle, the PMF at a given end-to-end distance should represent a thermodynamic average over all conformations compatible with that coordinate. Under ideal sampling conditions, structural substates sharing the same end-to-end distance would be integrated into a single free energy estimate. The sensitivity of the PMFs and ΔΔG° values to equilibration length therefore suggests that the two equilibration protocols are not sampling identical conformational ensembles at fixed end-to-end distance.

One possible explanation is the presence of slowly changing structural degrees of freedom that are only weakly coupled to the chosen reaction coordinate. While end-to-end distance captures the global extension of the hairpin and is attractive because it mimics optical tweezer experiments, (48) it does not explicitly describe base stacking, hydrogen-bonding patterns, loop conformations, or other structural features that may evolve on longer timescales. As a result, multiple metastable conformational substates may exist within a single umbrella window despite sharing similar end-to-end distances. Extended equilibration may permit relaxation into alternative substates that are sampled less frequently during shorter equilibration. The reduced hydrogen-bonding and stacking persistence observed in the 100 ns simulations is consistent with such a scenario, suggesting that longer equilibration promotes sampling of less structured conformational ensembles.

Importantly, our results do not demonstrate that the 2 ns protocol samples the correct thermodynamic ensemble while the 100 ns protocol does not. Rather, the data indicate that the structural ensembles sampled by the two protocols differ substantially and that these differences propagate directly into the calculated free energies. The observation that the 2 ns simulations exhibit better agreement with both experiment and nearest-neighbor-estimated free energy changes suggests that the corresponding ensembles may be more representative of the experimentally relevant folding landscape for these systems. We hypothesize that the better agreement observed with the 2 ns equilibration depends on metastability of native-like structures that are simply lost with longer equilibration. By the umbrella sampling window centered at 30 Å, in the intermediate ranges of end-to-end distances, 1600 ns of equilibration has already accumulated for the coordinates. Force fields are known to not accurately stabilize native RNA structures, (7,49–54) and we further hypothesize that the longer equilibration is revealing these inaccuracies. These simulations were performed with the conventional Amber OL3 force field. Future studies could test this using improved force fields that might not suffer from loss of accuracy with increased equilibration times. (55,56)

The sequence dependence of the equilibration effect further supports the view that hidden structural variables play an important role. GUGAAA and UUAAUU exhibited the strongest PMF flattening and among the largest equilibration-dependent changes in ΔΔG°, whereas CUGGGA was comparatively insensitive to equilibration length. GUAAUA displayed a distinct response characterized by a PMF reshaping rather than simple flattening. These observations indicate that equilibration length does not influence all RNA hairpins uniformly and that the accessibility of alternative conformational substates is likely sequence dependent.

## METHODS

### Umbrella Sampling Molecular Dynamics Simulation Protocol

The end-to-end distance corresponding to the distance between the 5’ and 3’ hydroxyl oxygens. The reaction coordinate was divided into series of overlapping windows with end-to-end distance ranging from 15 Å to 45 Å separated by 1 Å interval, by applying a harmonic restraint of force constant 3 kcal mol^-1^ Å^-1^ on the hydroxyl oxygen of the terminal residues. The simulations were performed using AMBER 20 and the RNA.OL3 force field with Li-Merz parameters for monovalent ions. (55–60) In each window, the negatively charged starting structure was neutralized by adding Na^+^ counterions and then solvated in OPC water (61) in an isometric truncated octahedron box with periodic boundary conditions and a 10 Å buffer. Additional Na^+^ and Cl^-^ ions were added to bring the molarity to 1 M. The protocol for addition of ions was in accordance to recommendations contained in Machado & Pantano. (62) The solvated system was minimized using 500 steps of steepest descent followed by an additional 500 steps of conjugate gradient minimization algorithms to bring the structures to the nearest minimum in the forcefield. The minimization was a two-stage process, with the first stage carried out by placing a weak harmonic restraint on the heavy atoms while allowing water and lighter atoms to minimize, followed by the removal of the restraint in the second minimization stage. The system was heated at NVT for 200 ps to 310 K, the temperature at which the experimental data for the free energy of stretching was obtained. Equilibration of the system was done at NPT for 2 or 100 ns.

Subsequent windows were solvated and equilibrated with starting structures being the last frame in the equilibration of the preceding window with the original solvent atoms stripped. Production molecular dynamics were run in each window for 600 ns. Each window was maintained at 310 K using a Langevin thermostat with a collision frequency of 2 ps^-1^ while a Monte Carlo barostat with a 100-step attempt frequency was used to regulate the pressure. (63,64) Hydrogen motion was constrained with SHAKE, allowing for a time step of 2 fs. (65)

### Free energy Calculation

For all four replicas of each system in both equilibration schemes, the end-to-end distance was calculated at each frame of their respective trajectories. We used the Grossfield implementation of the weighted histogram analysis method (WHAM) to assign relative probabilities to the conformational ensemble at discrete bins along the reaction coordinate. (25) A total of 311 bins ranging from 14.95 to 45.05 Å were specified as inputs to WHAM, in addition to a tolerance of 10^-6^ kcal/mol. The per-bin free energy values, derived by taking the natural logarithm of the relative probabilities, generated the potential of mean force (PMF) profiles. Free energy curves were plotted for individual replicates and for the pooled simulation generated by concatenating the trajectories of the four replicates of a system. The pooled PMF was used to represent the mean estimate of the free energy curve, while the replica-specific PMFs were retained to assess reproducibility across independent simulations. For an ergodic simulation, partition function for slack and stretched states can be defined at a specific cutoff, and estimated as the ratio of the sum of probabilities of bins below and above the cutoff, respectively (equation 1).

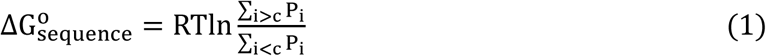

where Pi is the probability of the i^th^ bin, c is the cutoff, R is the molar gas constant in kcal/mol/K and T is the temperature in K.

### Validation of Predictions with Reference Data

We employed the thermodynamic cycle (Figure 3) to facilitate comparison between the free energy change of stretching and its estimate from nearest-neighbor (NN) rules and/or optical melting experiment. The necessity for the thermodynamic cycle rests on the disparity between thermal and force denaturation states, hence a more rational comparison must be done with respect to a pair of sequences rather than a single sequence. (34,37,46) We computed ΔΔG° stretch (equation 2) which is the relative free energy change between Sequence 1 and Sequence 2 consequent to force denaturation.

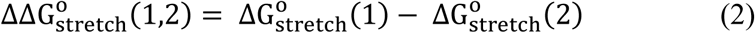

Similarly, the relative free energy from optical melting was calculated as follows:

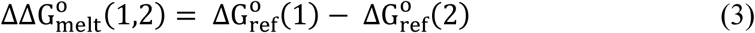

Where ΔG° is the reference optical melting data (or nearest neighbor-estimated stability) for the free energy change of a sequence.

Employing the thermodynamic cycle in Figure 3, the ΔΔG° from optical melting is:

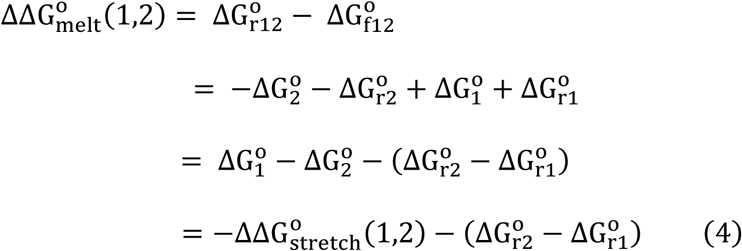

The last term in the parenthesis in equation 4 is often assumed to be sequence independent, (46) and therefore:

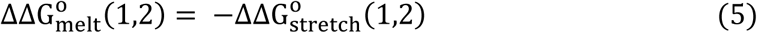

This provided the basis for which we validated simulated relative folding stability against reference data from optical melting or nearest neighbor rules.

### Uncertainty Estimation of **ΔG**°

The ΔG° resulting from pooling all replicas was taken as the mean and uncertainties were obtained from the standard error of the mean:

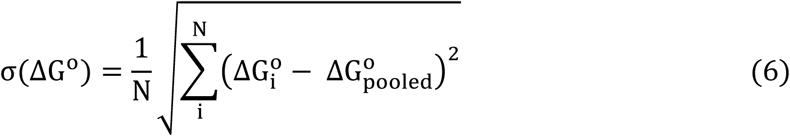

where N = 4, the number of replicates. The uncertainty on the estimation of ΔΔG^O^ for a pair of sequences (1,2) was obtained following the rules of error propagation upon addition or subtraction of quantities:

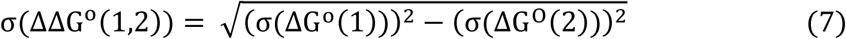

### Base-stacking and Hydrogen Bond analysis

Base-stacking interactions were quantified from the production trajectories for each hairpin, equilibration protocol, replicate, and umbrella window using CPPTRAJ. (66) Stacking was evaluated for all residue pairs *i*, *j*, where *i* < *j*, across the 12-residue hairpin. For each residue, a base plane was defined using the heavy atoms N1, N3, C2, C4, C5, and C6. For every residue pair, two quantities were calculated: the distance between the selected base atoms and the angle between the corresponding base-plane vectors. A residue pair was classified as stacked when the inter-base distance was less than 4.5 Å and the angle between base-plane vectors was less than 45°. (67) The stacking occupancy for each residue pair in each umbrella window was then computed as the fraction of trajectory frames satisfying both criteria.

Two stacking summaries were calculated. First, adjacent stacking was computed by summing stacking occupancies over neighboring residue pairs (*i*, *i* + 1). Second, total stacking was computed by summing stacking occupancies over all residue pairs (*i*, *j*), where *i* < *j*. These quantities were calculated separately for each of the four independent replicates and for each umbrella window. The mean and standard deviation across replicates were then reported as a function of end-to-end distance for the 2 ns and 100 ns equilibration protocols.

Hydrogen-bond analysis was also performed using CPPTRAJ. Hydrogen bonds were identified using a donor-acceptor distance cutoff of 3.5 Å and a donor-hydrogen-acceptor angle cutoff of 135°. For each umbrella window, the mean hydrogen-bond count was calculated across the four independent replicas. To characterize structural changes along the unfolding pathway, hydrogen bonds were separated into two categories: native hydrogen bonds present in the folded hairpin structure and non-neighboring base-base hydrogen bonds in simulation. The latter category was used to monitor the formation of alternative hydrogen-bonding interactions as native contacts were disrupted during extension.

### Optical Melting Experiments

Oligonucloetides were synthesized using β-cyanoethyl phosphoramidite chemistry on a BioAutomation MerMade12 DNA/RNA synthesizer with phosphoramidites from ChemPharma. (68) Deprotection was performed first with aqueous ammonia/ethanol (3/1 v/v) for 3 h at 55 °C and then treatment with triethylamine trihydrofluoride. Oligonucleotides were purified by thin layer chromatography with 1-propanol/aqueous ammonia/water (55/35/10 v/v/v) as the mobile phase. (44)

Optical melting experiments were performed using a JASCO V-650 UV/Vis spectrophotometer at 260 nm in a solvent of 1 M sodium chloride, 20 mM sodium cacodylate, and 0.5 mM Na2EDTA, pH 7.0. Strand concentrations were determined using extinction coefficients estimated from a nearest neighbor model and absorbances at 90 °C. (69,70) A series of experiments were performed at 7 concentrations from 0.3 mM to 7 μM. Curves were fit using MeltWin and reported ΔH°, ΔS°, and ΔG°37 are averages from the individual curve fits. (71)

## ACKNOWLEDGEMENTS

This work was supported by NIH grant R35GM145283 to D.H.M. and by National Science Center of Poland grants 2022/45/B/ST4/03586 to R.K. and 2021/41/B/NZ1/03819 and 2020/39/B/NZ1/03054 to E.K.

## Notes

### Competing Interest Statement

The authors have declared no competing interest.

## REFERENCES

1. Mattick, J. S., P. P. Amaral, P. Carninci, S. Carpenter, H. Y. Chang, L. L. Chen, R. Chen, C. Dean, M. E. Dinger, K. A. Fitzgerald, T. R. Gingeras, M. Guttman, T. Hirose, M. Huarte, R. Johnson, C. Kanduri, P. Kapranov, J. B. Lawrence, J. T. Lee, J. T. Mendell, T. R. Mercer, K. J. Moore, S. Nakagawa, J. L. Rinn, D. L. Spector, I. Ulitsky, Y. Wan, J. E. Wilusz, and M. Wu. 2023. Long non-coding RNAs: definitions, functions, challenges and recommendations. Nat Rev Mol Cell Biol. 24(6):430–447, doi: 10.1038/s41580-022-00566-8, https://www.ncbi.nlm.nih.gov/pubmed/36596869.

2. Eddy, S. R. 2001. Non-coding RNA genes and the modern RNA world. Nat Rev Genet. 2(12):919–929, doi: 10.1038/35103511, http://www.ncbi.nlm.nih.gov/pubmed/11733745.

3. Hombach, S., and M. Kretz. 2016. Non-coding RNAs: Classification, Biology and Functioning. Adv Exp Med Biol. 937:3–17, doi: 10.1007/978-3-319-42059-2_1, https://www.ncbi.nlm.nih.gov/pubmed/27573892.

4. Korostelev, A., S. Trakhanov, M. Laurberg, and H. F. Noller. 2006. Crystal structure of a 70S ribosome-tRNA complex reveals functional interactions and rearrangements. Cell. 126(6):1065–1077, doi: S0092-8674(06)01146-9, http://www.ncbi.nlm.nih.gov/pubmed/16962654.

5. Schroeder, G. M., O. Akinyemi, J. Malik, C. Focht, E. M. Pritchett, C. D. Baker, J. P. McSally, J. L. Jenkins, D. H. Mathews, and J. E. Wedekind. 2023. A riboswitch separated from its ribosome-binding site still regulates translation. Nucleic Acids Res. 51(5):2464–2484, doi: 10.1093/nar/gkad056.

6. Srivastava, Y., O. Akinyemi, T. C. Rohe, E. M. Pritchett, C. D. Baker, A. Sharma, J. L. Jenkins, D. H. Mathews, and J. E. Wedekind. 2024. Two riboswitch classes that share a common ligand-binding fold show major differences in the ability to accommodate mutations. Nucleic Acids Res. 52(21):13152–13173, doi: 10.1093/nar/gkae886, https://www.ncbi.nlm.nih.gov/pubmed/39413212.

7. Akinyemi, O., S. D. Kennedy, J. P. McSally, and D. H. Mathews. 2026. NMR and molecular dynamics demonstrate the RNA internal loop GAGU is dynamic and adjacent basepairs determine conformational preference. Biophys J. 125(1):204–217, doi: 10.1016/j.bpj.2025.11.026, https://www.ncbi.nlm.nih.gov/pubmed/41261770.

8. Hammond, N. B., B. S. Tolbert, R. Kierzek, D. H. Turner, and S. D. Kennedy. 2010. RNA internal loops with tandem AG pairs: the structure of the 5’GAGU/3’UGAG loop can be dramatically different from others, including 5’AAGU/3’UGAA. Biochemistry. 49(27):5817–5827, doi: 10.1021/bi100332r, http://www.ncbi.nlm.nih.gov/pubmed/20481618.

9. Zhang, H., L. Zhang, D. H. Mathews, and L. Huang. 2020. LinearPartition: linear-time approximation of RNA folding partition function and base-pairing probabilities. Bioinformatics. 36(Supplement_1):i258–i267, doi: 10.1093/bioinformatics/btaa460, http://www.ncbi.nlm.nih.gov/pubmed/32657379.

10. Zuber, J., S. J. Schroeder, H. Sun, D. H. Turner, and D. H. Mathews. 2022. Nearest neighbor rules for RNA helix folding thermodynamics: improved end effects. Nucleic Acids Res. 50(9):5251–5262, doi: 10.1093/nar/gkac261, https://www.ncbi.nlm.nih.gov/pubmed/35524574.

11. Jouravleva, K., J. Vega-Badillo, and P. D. Zamore. 2022. Principles and pitfalls of high-throughput analysis of microRNA-binding thermodynamics and kinetics by RNA Bind-n-Seq. Cell Rep Methods. 2(3):100185, doi: 10.1016/j.crmeth.2022.100185, https://www.ncbi.nlm.nih.gov/pubmed/35475222.

12. Levintov, L., S. Paul, and H. Vashisth. 2021. Reaction Coordinate and Thermodynamics of Base Flipping in RNA. J Chem Theory Comput. 17(3):1914–1921, doi: 10.1021/acs.jctc.0c01199, https://www.ncbi.nlm.nih.gov/pubmed/33594886.

13. Mandic, A., R. L. Hayes, H. Lammert, R. R. Cheng, and J. N. Onuchic. 2019. Structure-Based Model of RNA Pseudoknot Captures Magnesium-Dependent Folding Thermodynamics. J Phys Chem B. 123(7):1505–1511, doi: 10.1021/acs.jpcb.8b10791, https://www.ncbi.nlm.nih.gov/pubmed/30676755.

14. Joseph, J. A., J. R. Espinosa, I. Sanchez-Burgos, A. Garaizar, D. Frenkel, and R. Collepardo-Guevara. 2021. Thermodynamics and kinetics of phase separation of protein-RNA mixtures by a minimal model. Biophys J. 120(7):1219–1230, doi: 10.1016/j.bpj.2021.01.031, https://www.ncbi.nlm.nih.gov/pubmed/33571491.

15. Banas, P., D. Hollas, M. Zgarbova, P. Jurecka, M. Orozco, T. E. Cheatham, J. Sponer, and M. Otyepka. 2010. Performance of molecular mechanics force fields for RNA simulations: Stability of UUCG and GNRA hairpins. J. Chem. Theory Comput. 6:3836–3849.

16. Mrazikova, K., V. Mlynsky, P. Kuhrova, P. Pokorna, H. Kruse, M. Krepl, M. Otyepka, P. Banas, and J. Sponer. 2020. UUCG RNA Tetraloop as a Formidable Force-Field Challenge for MD Simulations. J Chem Theory Comput. 16(12):7601–7617, doi: 10.1021/acs.jctc.0c00801, http://www.ncbi.nlm.nih.gov/pubmed/33215915.

17. Barducci, A., G. Bussi, and M. Parrinello. 2008. Well-tempered metadynamics: a smoothly converging and tunable free-energy method. Phys Rev Lett. 100(2):020603, doi: 10.1103/PhysRevLett.100.020603, https://www.ncbi.nlm.nih.gov/pubmed/18232845.

18. Roy, R., A. Mishra, S. Poddar, D. Nayak, and P. Kar. 2022. Investigating the mechanism of recognition and structural dynamics of nucleoprotein-RNA complex from Peste des petits ruminants virus via Gaussian accelerated molecular dynamics simulations. J Biomol Struct Dyn. 40(5):2302–2315, doi: 10.1080/07391102.2020.1838327, https://www.ncbi.nlm.nih.gov/pubmed/33089759.

19. Liu, X., and C. L. Brooks Iii. 2024. Enhanced Sampling of Buried Charges in Free Energy Calculations Using Replica Exchange with Charge Tempering. J Chem Theory Comput. 20(3):1051–1061, doi: 10.1021/acs.jctc.3c00993, https://www.ncbi.nlm.nih.gov/pubmed/38232295.

20. Hasse, T., and Y. M. Huang. 2024. Multiple Parameter Replica Exchange Gaussian Accelerated Molecular Dynamics for Enhanced Sampling and Free Energy Calculation of Biomolecular Systems. J Chem Theory Comput. 20(15):6485–6499, doi: 10.1021/acs.jctc.4c00501, https://www.ncbi.nlm.nih.gov/pubmed/39085770.

21. Govind Kumar, V., A. Polasa, S. Agrawal, T. K. S. Kumar, and M. Moradi. 2023. Binding affinity estimation from restrained umbrella sampling simulations. Nat Comput Sci. 3(1):59–70, doi: 10.1038/s43588-022-00389-9, https://www.ncbi.nlm.nih.gov/pubmed/38177953.

22. Yildirim, I., H. Park, M. D. Disney, and G. C. Schatz. 2013. A dynamic structural model of expanded RNA CAG repeats: a refined X-ray structure and computational investigations using molecular dynamics and umbrella sampling simulations. J Am Chem Soc. 135(9):3528–3538, doi: 10.1021/ja3108627, https://www.ncbi.nlm.nih.gov/pubmed/23441937.

23. Dickson, A., M. Maienschein-Cline, A. Tovo-Dwyer, J. R. Hammond, and A. R. Dinner. 2011. Flow-Dependent Unfolding and Refolding of an RNA by Nonequilibrium Umbrella Sampling. J Chem Theory Comput. 7(9):2710–2720, doi: 10.1021/ct200371n, https://www.ncbi.nlm.nih.gov/pubmed/26605464.

24. Volynets, G. P., O. I. Gudzera, M. O. Usenko, O. B. Gorbatiuk, V. G. Bdzhola, I. M. Kotey, A. O. Balanda, A. O. Prykhod’ko, S. S. Lukashov, O. A. Chuk, O. I. Skydanovych, G. D. Yaremchuk, S. M. Yarmoluk, and M. A. Tukalo. 2025. Probing the Molecular Basis of Aminoacyl-Adenylate Affinity With Mycobacterium tuberculosis Leucyl-tRNA Synthetase Employing Molecular Dynamics, Umbrella Sampling Simulations and Site-Directed Mutagenesis. J Mol Recognit. 38(2):e3110, doi: 10.1002/jmr.3110, https://www.ncbi.nlm.nih.gov/pubmed/39478352.

25. Grossfield, A. WHAM: an implementation of the weighted histogram analysis method. doi: https://github.com/agrossfield/wham.

26. Ferguson, A. L. 2017. BayesWHAM: A Bayesian approach for free energy estimation, reweighting, and uncertainty quantification in the weighted histogram analysis method. J Comput Chem. 38(18):1583–1605, doi: 10.1002/jcc.24800, https://www.ncbi.nlm.nih.gov/pubmed/28475830.

27. Kim, J., T. Keyes, and J. E. Straub. 2011. Communication: Iteration-free, weighted histogram analysis method in terms of intensive variables. J Chem Phys. 135(6):061103, doi: 10.1063/1.3626150, https://www.ncbi.nlm.nih.gov/pubmed/21842919.

28. Yang, M., and A. D. MacKerell, Jr. 2015. Conformational sampling of oligosaccharides using Hamiltonian replica exchange with two-dimensional dihedral biasing potentials and the weighted histogram analysis method (WHAM). J Chem Theory Comput. 11(2):788–799, doi: 10.1021/ct500993h, https://www.ncbi.nlm.nih.gov/pubmed/25705140.

29. Zhu, F., and G. Hummer. 2012. Convergence and error estimation in free energy calculations using the weighted histogram analysis method. J Comput Chem. 33(4):453–465, doi: 10.1002/jcc.21989, https://www.ncbi.nlm.nih.gov/pubmed/22109354.

30. Ngo, V. A., I. Kim, T. W. Allen, and S. Y. Noskov. 2016. Estimation of Potentials of Mean Force from Nonequilibrium Pulling Simulations Using Both Minh-Adib Estimator and Weighted Histogram Analysis Method. J Chem Theory Comput. 12(3):1000–1010, doi: 10.1021/acs.jctc.5b01050, https://www.ncbi.nlm.nih.gov/pubmed/26799775.

31. Mathath, A. V., B. K. Das, and D. Chakraborty. 2023. Designing Reaction Coordinate for Ion-Induced Pore-Assisted Mechanism of Halide Ions Permeation through Lipid Bilayer by Umbrella Sampling. J Chem Inf Model. 63(24):7778–7790, doi: 10.1021/acs.jcim.3c01683, https://www.ncbi.nlm.nih.gov/pubmed/38050816.

32. Li, G., Z. Li, L. Gao, S. Chen, G. Wang, and S. Li. 2023. Combined molecular dynamics and coordinate driving method for automatically searching complicated reaction pathways. Phys Chem Chem Phys. 25(35):23696–23707, doi: 10.1039/d3cp02443a, https://www.ncbi.nlm.nih.gov/pubmed/37610711.

33. Cheng, T., W. Ma, H. Luo, Y. Ye, and K. Yan. 2022. Manipulating Reaction Energy Coordinate Landscape of Mechanochemical Diaza-Cope Rearrangement. Molecules. 27(8), doi: 10.3390/molecules27082570, https://www.ncbi.nlm.nih.gov/pubmed/35458767.

34. Smith, L. G., Z. Tan, A. Spasic, D. Dutta, L. A. Salas-Estrada, A. Grossfield, and D. H. Mathews. 2018. Chemically Accurate Relative Folding Stability of RNA Hairpins from Molecular Simulations. J Chem Theory Comput. doi: 10.1021/acs.jctc.8b00633, http://www.ncbi.nlm.nih.gov/pubmed/30375860.

35. Morosyuk, S. V., P. R. Cunningham, and J. SantaLucia, Jr. 2001. Structure and function of the conserved 690 hairpin in Escherichia coli 16 S ribosomal RNA. II. NMR solution structure. J Mol Biol. 307(1):197–211, doi: 10.1006/jmbi.2000.4431, https://www.ncbi.nlm.nih.gov/pubmed/11243814.

36. Brodsky, A. S., and J. R. Williamson. 1997. Solution structure of the HIV-2 TAR-argininamide complex. J Mol Biol. 267(3):624–639, doi: 10.1006/jmbi.1996.0879, https://www.ncbi.nlm.nih.gov/pubmed/9126842.

37. Spasic, A., J. Serafini, and D. H. Mathews. 2012. The Amber ff99 Force Field Predicts Relative Free Energy Changes for RNA Helix Formation. J Chem Theory Comput. 8(7):2497–2505, doi: 10.1021/ct300240k, https://www.ncbi.nlm.nih.gov/pubmed/23112748.

38. Fountain, M. A., M. J. Serra, T. R. Krugh, and D. H. Turner. 1996. Structural features of a six-nucleotide RNA hairpin loop found in ribosomal RNA. Biochemistry. 35(21):6539–6548, doi: 10.1021/bi952697k, http://www.ncbi.nlm.nih.gov/pubmed/8639602.

39. Arora, A., and S. Maiti. 2009. Differential biophysical behavior of human telomeric RNA and DNA quadruplex. J Phys Chem B. 113(30):10515–10520, doi: 10.1021/jp810638n, https://www.ncbi.nlm.nih.gov/pubmed/19572668.

40. Pandey, S., P. Agarwala, and S. Maiti. 2013. Effect of loops and G-quartets on the stability of RNA G-quadruplexes. J Phys Chem B. 117(23):6896–6905, doi: 10.1021/jp401739m, https://www.ncbi.nlm.nih.gov/pubmed/23683360.

41. Mergny, J. L., and L. Lacroix. 2009. UV Melting of G-Quadruplexes. Curr Protoc Nucleic Acid Chem. Chapter 17:17 11 11-17 11 15, doi: 10.1002/0471142700.nc1701s37, https://www.ncbi.nlm.nih.gov/pubmed/19488970.

42. Mittal, A., D. H. Turner, and D. H. Mathews. 2024. NNDB: An Expanded Database of Nearest Neighbor Parameters for Predicting Stability of Nucleic Acid Secondary Structures. J Mol Biol. 436(17):168549, doi: 10.1016/j.jmb.2024.168549, https://www.ncbi.nlm.nih.gov/pubmed/38522645.

43. Mathews, D. H., J. Sabina, M. Zuker, and D. H. Turner. 1999. Expanded sequence dependence of thermodynamic parameters improves prediction of RNA secondary structure. J Mol Biol. 288(5):911–940, doi: 10.1006/jmbi.1999.2700, http://www.ncbi.nlm.nih.gov/pubmed/10329189.

44. Xia, T., J. SantaLucia, Jr., M. E. Burkard, R. Kierzek, S. J. Schroeder, X. Jiao, C. Cox, and D. H. Turner. 1998. Thermodynamic parameters for an expanded nearest-neighbor model for formation of RNA duplexes with Watson-Crick base pairs. Biochemistry. 37(42):14719–14735, doi: 10.1021/bi9809425, http://www.ncbi.nlm.nih.gov/pubmed/9778347.

45. Zuber, J., B. J. Cabral, I. McFadyen, D. M. Mauger, and D. H. Mathews. 2018. Analysis of RNA nearest neighbor parameters reveals interdependencies and quantifies the uncertainty in RNA secondary structure prediction. RNA. 24(11):1568–1582, doi: 10.1261/rna.065102.117, https://www.ncbi.nlm.nih.gov/pubmed/30104207.

46. Tinoco, I., Jr., P. T. Li, and C. Bustamante. 2006. Determination of thermodynamics and kinetics of RNA reactions by force. Q Rev Biophys. 39(4):325–360, doi: 10.1017/S0033583506004446, https://www.ncbi.nlm.nih.gov/pubmed/17040613.

47. Smith, L. G., J. Zhao, D. H. Mathews, and D. H. Turner. 2017. Physics-based all-atom modeling of RNA energetics and structure. WIREs RNA. 8, e1422(5), doi: 10.1002/wrna.1422, http://www.ncbi.nlm.nih.gov/pubmed/28815951.

48. Stephenson, W., S. Keller, R. Santiago, J. E. Albrecht, P. N. Asare-Okai, S. A. Tenenbaum, M. Zuker, and P. T. Li. 2014. Combining temperature and force to study folding of an RNA hairpin. Phys Chem Chem Phys. 16(3):906–917, doi: 10.1039/c3cp52042k, https://www.ncbi.nlm.nih.gov/pubmed/24276015.

49. Bergonzo, C., N. M. Henriksen, D. R. Roe, and T. E. Cheatham, 3rd. 2015. Highly sampled tetranucleotide and tetraloop motifs enable evaluation of common RNA force fields. RNA. 21(9):1578–1590, doi: 10.1261/rna.051102.115, http://www.ncbi.nlm.nih.gov/pubmed/26124199.

50. Bergonzo, C., and T. E. Cheatham, 3rd. 2015. Improved Force Field Parameters Lead to a Better Description of RNA Structure. J Chem Theory Comput. 11(9):3969–3972, doi: 10.1021/acs.jctc.5b00444, http://www.ncbi.nlm.nih.gov/pubmed/26575892.

51. Mlynsky, V., P. Kuhrova, M. Pykal, M. Krepl, P. Stadlbauer, M. Otyepka, P. Banas, and J. Sponer. 2025. Can We Ever Develop an Ideal RNA Force Field? Lessons Learned from Simulations of the UUCG RNA Tetraloop and Other Systems. J Chem Theory Comput. 21(8):4183–4202, doi: 10.1021/acs.jctc.4c01357, https://www.ncbi.nlm.nih.gov/pubmed/39813107.

52. Bottaro, S., P. Banas, J. Sponer, and G. Bussi. 2016. Free Energy Landscape of GAGA and UUCG RNA Tetraloops. J Phys Chem Lett. 7(20):4032–4038, doi: 10.1021/acs.jpclett.6b01905, http://www.ncbi.nlm.nih.gov/pubmed/27661094.

53. Zgarbova, M., P. Jurecka, P. Banas, M. Havrila, J. Sponer, and M. Otyepka. 2017. Noncanonical alpha/gamma Backbone Conformations in RNA and the Accuracy of Their Description by the AMBER Force Field. J Phys Chem B. 121(11):2420–2433, doi: 10.1021/acs.jpcb.7b00262, https://www.ncbi.nlm.nih.gov/pubmed/28290207.

54. Yildirim, I., H. A. Stern, J. D. Tubbs, S. D. Kennedy, and D. H. Turner. 2011. Benchmarking AMBER force fields for RNA: comparisons to NMR spectra for single-stranded r(GACC) are improved by revised chi torsions. J Phys Chem B. 115(29):9261–9270, doi: 10.1021/jp2016006, http://www.ncbi.nlm.nih.gov/pubmed/21721539.

55. Zgarbova, M., M. Otyepka, J. Sponer, A. Mladek, P. Banas, T. E. Cheatham, 3rd, and P. Jurecka. 2011. Refinement of the Cornell et al. Nucleic Acids Force Field Based on Reference Quantum Chemical Calculations of Glycosidic Torsion Profiles. J Chem Theory Comput. 7(9):2886–2902, doi: 10.1021/ct200162x, https://www.ncbi.nlm.nih.gov/pubmed/21921995.

56. Perez, A., I. Marchan, D. Svozil, J. Sponer, T. E. Cheatham, 3rd, C. A. Laughton, and M. Orozco. 2007. Refinement of the AMBER force field for nucleic acids: improving the description of alpha/gamma conformers. Biophys J. 92(11):3817–3829, doi: 10.1529/biophysj.106.097782, https://www.ncbi.nlm.nih.gov/pubmed/17351000.

57. Wang, J., P. Cieplak, and P. A. Kollman. 2000. How well does a restrained electrostatic potential (RESP) model perform in calculating conformational energies of organic and biological molecules? J Comput Chem. 21:1049–1074.

58. Sengupta, A., Z. Li, L. F. Song, P. Li, and K. M. Merz, Jr. 2021. Parameterization of Monovalent Ions for the OPC3, OPC, TIP3P-FB, and TIP4P-FB Water Models. J Chem Inf Model. 61(2):869–880, doi: 10.1021/acs.jcim.0c01390, https://www.ncbi.nlm.nih.gov/pubmed/33538599.

59. Li, P., L. F. Song, and K. M. Merz, Jr. 2015. Systematic Parameterization of Monovalent Ions Employing the Nonbonded Model. J Chem Theory Comput. 11(4):1645–1657, doi: 10.1021/ct500918t, https://www.ncbi.nlm.nih.gov/pubmed/26574374.

60. Case, D. A., T. E. Cheatham, 3rd, T. Darden, H. Gohlke, R. Luo, K. M. Merz, Jr., A. Onufriev, C. Simmerling, B. Wang, and R. J. Woods. 2005. The Amber biomolecular simulation programs. J Comput Chem. 26(16):1668–1688, doi: 10.1002/jcc.20290, https://www.ncbi.nlm.nih.gov/pubmed/16200636.

61. Izadi, S., R. Anandakrishnan, and A. V. Onufriev. 2014. Building Water Models: A Different Approach. J Phys Chem Lett. 5(21):3863–3871, doi: 10.1021/jz501780a, https://www.ncbi.nlm.nih.gov/pubmed/25400877.

62. Machado, M. R., and S. Pantano. 2020. Split the Charge Difference in Two! A Rule of Thumb for Adding Proper Amounts of Ions in MD Simulations. J Chem Theory Comput. 16(3):1367–1372, doi: 10.1021/acs.jctc.9b00953, https://www.ncbi.nlm.nih.gov/pubmed/31999456.

63. Loncharich, R. J., B. R. Brooks, and R. W. Pastor. 1992. Langevin dynamics of peptides: the frictional dependence of isomerization rates of N-acetylalanyl-N’-methylamide. Biopolymers. 32(5):523–535, doi: 10.1002/bip.360320508, https://www.ncbi.nlm.nih.gov/pubmed/1515543.

64. Gomez, Y. K., A. M. Natale, J. Lincoff, C. W. Wolgemuth, J. M. Rosenberg, and M. Grabe. 2022. Taking the Monte-Carlo gamble: How not to buckle under the pressure! J Comput Chem. 43(6):431–434, doi: 10.1002/jcc.26798, https://www.ncbi.nlm.nih.gov/pubmed/34921560.

65. Ryckaert, J. P., G. Ciccotti, and H. J. C. Berendsen. 1977. Numerical integration of the cartesian equations of motion of a system with constraints: molecular dynamics of n-alkanes. J Comput Phys. 23:327–341, doi: 10.1016/0021-9991(77)90098-5.

66. Roe, D. R., and T. E. Cheatham 3rd,. 2013. PTRAJ and CPPTRAJ: Software for Processing and Analysis of Molecular Dynamics Trajectory Data. J Chem Theory Comput. 9(7):3084–3095, doi: 10.1021/ct400341p, https://www.ncbi.nlm.nih.gov/pubmed/26583988.

67. Taghavi, A., I. Riveros, D. J. Wales, and I. Yildirim. 2022. Evaluating Geometric Definitions of Stacking for RNA Dinucleoside Monophosphates Using Molecular Mechanics Calculations. J Chem Theory Comput. 18(6):3637–3653, doi: 10.1021/acs.jctc.2c00178, https://www.ncbi.nlm.nih.gov/pubmed/35652685.

68. Beaucage, S. L., and M. H. Caruthers. 1981. Deoxynucleoside phosphoramidites - A new class of key intermediates for deoxypolynucleotide synthesis. Tetrahedron Letters. 22(20):1859–1862, doi: 10.1016/S0040-4039(01)90461-7.

69. Borer, P. N. 1975. Optical properties of nucleic acids, absorption and circular dichroism spectra. In Handbook of Biochemistry and Molecular Biology: Nucleic Acids, 3rd Ed. G. D. Fasman, editor. CRC Press, Cleveland, OH, pp. 589–595.

70. Richards, E. G. 1975. Use of tables in calculation of absorption, optical rotatory dispersion and circular dichroism of polyribonucleotides. In Handbook of Biochemistry and Molecular Biology: Nucleic Acids, 3rd Ed. G. D. Fasman, editor. CRC Press, Cleveland, OH, pp. 596–603.

71. McDowell, J. A., L. He, X. Chen, and D. H. Turner. 1997. Investigation of the structural basis for thermodynamic stabilities of tandem GU wobble pairs: NMR structures of (rGGAGUUCC)2 and (rGGAUGUCC)2. Biochemistry. 36(26):8030–8038, doi: 10.1021/bi970122c, http://www.ncbi.nlm.nih.gov/pubmed/9201950.

